# Different sensory information is used for state estimation when stationary or moving

**DOI:** 10.1101/2023.09.01.555979

**Authors:** Aaron L Wong, Alyssa N Eyssalenne, Luke Carter, Amanda S Therrien

## Abstract

The accurate estimation of limb state is necessary for movement planning and execution. While state estimation requires both feedforward and feedback information, we focus here on the latter. Prior literature has shown that integrating visual and proprioceptive feedback improves estimates of static limb position. However, differences in visual and proprioceptive feedback delays suggest that multisensory integration could be disadvantageous when the limb is moving. We formalized this hypothesis by modeling feedback-based state estimation using the longstanding maximum likelihood estimation model of multisensory integration, which we updated to account for sensory delays. Our model predicted that the benefit of multisensory integration was largely lost when the limb was passively moving. We tested this hypothesis in a series of experiments in human subjects that compared the degree of interference created by discrepant visual or proprioceptive feedback when estimating limb position either statically at the end of the movement or dynamically at movement midpoint. In the static case, we observed significant interference: discrepant feedback in one modality systematically biased sensory estimates based on the other modality. However, no interference was seen in the dynamic case: participants could ignore sensory feedback from one modality and accurately reproduce the motion indicated by the other modality. Together, these findings suggest that the sensory feedback used to compute a state estimate differs depending on whether the limb is stationary or moving. While the former may tend toward multimodal integration, the latter is more likely to be based on feedback from a single sensory modality.

**SIGNIFICANCE STATEMENT:** The estimation of limb state involves feedforward and feedback information. While estimation based on feedback has been well studied when the limb is stationary, it is unknown if similar sensory processing supports limb position estimates when moving. Using a computational model and behavioral experiments, we show that feedback-based state estimation may involve multisensory integration in the static case, but it is likely based on a single modality when the limb is moving. We suggest that this difference may stem from visual and proprioceptive feedback delays.

## INTRODUCTION

Knowing the current state of the limb (i.e., its position and motion) is critical for the planning and online control of movement. Although limb state estimation is thought to involve both feedforward and feedback processes (Stengel, 1994; Wolpert et al., 1995), here we focus on the latter case. Previous work studying feedback integration in state estimation has shown that it is advantageous to integrate information from multiple sensory inputs to reduce overall uncertainty and produce a more reliable position estimate (Berniker and Kording, 2011; Ernst and Banks, 2002; van Beers et al., 1999, 1996). Multisensory integration is evident when the limb states sensed by two modalities (e.g., vision and proprioception) are in significant disagreement. The discrepancy is reduced by changing the relative reliance on vision versus proprioception (i.e., reweighting; Smeets et al., 2006) or by recalibrating one sense to align with the other (i.e., realignment; Block and Bastian, 2010; 2011, Rossi et al., 2021), resulting in a systematic biasing of sensory estimates in the direction that minimizes intermodal misalignment. An extreme example of sensory discrepancy reduction is the rubber hand illusion (Botvinick and Cohen, 1998), in which people experience the transient illusion that their proprioceptively sensed arm position is aligned with the visually observed location of a fake stationary arm. This illusion is induced by integrating synchronous visual and somatosensory feedback (i.e., simultaneous stroking of the fake and real arm).

Prior work studying multisensory integration has focused on state estimation when the limb is stationary. However, multisensory integration becomes problematic when the limb is moving, especially when one considers differences in the timing of feedback delays across modalities (Cluff et al., 2015; Crevecoeur et al., 2016; Kasuga et al., 2022). For example, in the macaque, there is a delay of approximately ∼70 ms for visual information to reach V1, whereas the delay of proprioceptive information to S1 is only ∼20 ms (Bair et al., 2002; Nowak et al., 1995; Raiguel et al., 1989; Song and Francis, 2013). In this context, multisensory integration requires the system to wait for feedback information to become available from the slowest modality or to integrate feedback from different modalities that are delayed to different extents. The former would lead to a state estimate that is chronically out-of-date, while the latter would increase estimation error and uncertainty since each sensory system would be reporting the position of the limb when it is in a different physical location. Either way, a feedback-based estimate of the moving limb that relies on multisensory integration is likely to be unreliable relative to an estimate made with the same sensory input when the limb is stationary. During active limb movements, this problem may be partially resolved by the availability of sensory prediction signals, which provide a real-time prediction of the current limb position and thus can be used to mitigate erroneous motor commands (Crevecoeur et al., 2016; Kasuga et al., 2022).

However, the availability of sensory prediction signals does not fully resolve the problem because real albeit delayed sensory feedback is still required to respond to unexpected environmental perturbations (Cluff et al., 2015). While it is possible that the benefits of multisensory integration outweigh the potential uncertainty introduced by moving for both active and passive movements, in some circumstances, it might be better to rely on a single source of sensory feedback.

We start from the perspective of updating the classic maximum likelihood estimation model of multisensory integration (Ernst and Banks, 2002; van Beers et al., 1996) to take into account differences in sensory delays. This updated model suggests that multisensory integration is advantageous when the limb is stationary or moving at very low speeds. However, this advantage appears to be lost when limb speed increases; in this case, the use of feedback from a single sensory modality may be more advantageous. To test this prediction of our model, we compared feedback integration in state estimation when the limb was stationary to when it was being passively moved and examined the extent to which estimates were dependent upon multisensory integration in these two conditions. We opted to focus on passive movements to study the sensory feedback integration process in the absence of any sensory prediction signals. We asked individuals to report a position or motion cued by one sensory modality (i.e., vision or proprioception) while ignoring discrepant feedback from a second modality. When the limb was stationary, we found that individuals exhibited a bias toward the discrepant sensory modality, suggesting that they relied on multisensory integration to compute state estimates. This finding is in line with prior literature. Individuals exhibited no bias when moving, however, suggesting that they were computing state estimates based largely on a single sensory modality. Across two follow-up experiments, we show that this difference in bias when the limb is stationary versus moving persists regardless of how the multisensory discrepancy is presented. Overall, our findings support the predictions of our computational model and indicate that feedback integration in state estimation differs when moving versus when stationary.

## MATERIALS AND METHODS

### Modeling

Multisensory integration for state estimation has previously been modeled using a maximum likelihood estimation (MLE) approach (Ernst and Banks, 2002). We propose a simple update to this approach to account for the effect of sensory delays during movement (see Fig. 5A). We begin with the assumption that the hand is moving at speed, 𝜐, and the temporal delay for feedback from sensory system, *i* (either Vision, V, or Proprioception, P), is 𝜏_𝑖_. For simplicity, we assume this model reflects a snapshot in time; that is, none of our parameters are time-dependent (or equivalently, all our parameters reflect the same instant in time). If we assume that sensory feedback is distributed as a Gaussian with a mean, 𝜇_𝑖_, and variance, 𝜎_𝑖_^2^, then the sensed position available at time, *t,* can be described as the position sensed 𝜏_𝑖_ seconds ago that is spatially displaced by an amount equal to the product of the sensory time delay and the movement speed:

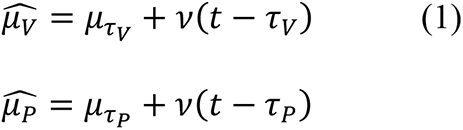

According to the MLE estimate, combining sensory feedback from the two modalities (e.g., akin to the bimodal condition in our task) should result in a Gaussian defined by:

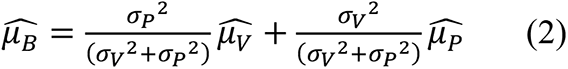

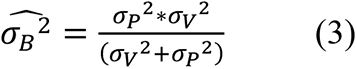

However, the unimodal sensory feedback that is actually available is the delayed sensory feedback given in the pair of equations (1) above. Hence, the actual integrated sensory feedback at time, *t,* is:

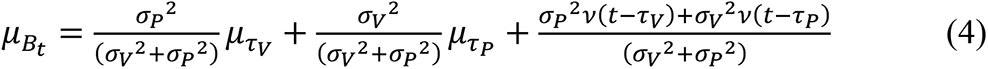

where the third term reflects the offset of the state estimate due to sensory delays. With this information, we can calculate the degree of overlap between the actual (equation 4) and ideal (equation 2) bimodal-estimate Gaussians (where the ideal bimodal estimate is calculated assuming that sensory feedback is available instantaneously, equal to equation (4) without the offset term). This overlap is equal to the intersection (area under both curves) of the actual and ideal bimodal-estimate Gaussians, which can be calculated by finding the point of intersection of the two Gaussians, *c*, summing the probability, P[Xactual > *c*], and the probability, P[Xideal < *c*], and normalizing by the union (total area) of both probability distributions. The overlap enables us to estimate the probability with which the two sensory estimates (actual and ideal) will correspond to one another. That is, the overlap between 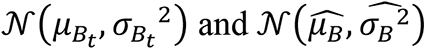 reflects the probability that the bimodal state estimate, accounting for sensory delays and movement speed, will fall within the ideal bimodal distribution had the limb been stationary. This probability tells us the degree to which an individual could expect to calculate a reasonable bimodal state estimate given their current (instantaneous) movement speed and the uncertainty associated with each sensory modality. Using this model, we calculated this probability over a range of physiologically reasonable movement speeds, sensory delays, and sensory variances to examine how these parameters change the degree to which an individual should rely on unimodal versus bimodal state estimates.

#### Experiments

A total of sixty-four right-handed adult neurotypical individuals were recruited for this study. Twenty-five individuals participated in Experiment 1, which consisted of two sessions spaced at minimum one week apart (average 17 days). Of the twenty-five individuals, twenty-two participated in both the static and dynamic estimation tasks (age 19-36 years, average 26.0 years, standard deviation 4.9 years; 17 identified as female, 4 identified as male, 1 preferred not to respond), one participated only in the static estimation task (25 year old who identified as female), and one participated only in the dynamic estimation task (29 year old who identified as male). One individual was excluded from Experiment 1 for failing to comply with task instructions. Twenty individuals participated in Experiment 2 (age 19-35 years, average 23.3 years, standard deviation 4.7 years; 15 identified as female, 5 identified as male). Nineteen individuals participated in Experiment 3. Of these nineteen participants, four were excluded for inability to follow directions; thus, we included data from 15 participants (age 20-33 years, average 25.1 years, standard deviation 3.2 years; 10 identified as female, 5 identified as male). Sample sizes for each experiment were chosen to be comparable to previous studies utilizing a similar task (e.g., Block and Bastian, 2010; Block and Liu, 2023; Reuschel et al., 2010). All participants were naïve to the purposes of this study and provided written informed consent. Experimental methods were approved by the Thomas Jefferson University Institutional Review Board, and participants were compensated for their participation at a fixed hourly payment ($20/hr).

Participants were seated in a Kinarm Exoskeleton Lab (Kinarm, Kingston, ON, Canada), which restricted motion of the arms to the horizontal plane. Hands, forearms, and upper arms were supported in troughs appropriately sized to each participant’s arms, and the linkage lengths of the robot were adjusted to match the limb segment lengths for each participant to allow smooth motion at the elbow and shoulder. Vision of the arm was obstructed by a horizontal mirror, through which participants could be shown targets and cursors in a veridical horizontal plane via an LCD monitor (60 Hz). Movement of the arm was recorded at 1000 Hz. For simplicity, we henceforth refer to the medial-lateral direction (i.e., movement along the x-axis) as “horizontal” and the anterior-posterior direction (i.e., movement along the y-axis) as “vertical”.

#### Experiment 1

In separate sessions with order counterbalanced, participants were asked to complete a static and a dynamic estimation task (Fig 1). Simulink code to run the tasks on the Kinarm is available on the lab GitHub page (https://github.com/CML-lab/KINARM_Static_Sensory_Integration and https://github.com/CML-lab/KINARM_Dynamic_Sensory_Integration respectively). Participants began each trial with the robot bringing their hand into one of five possible starting positions randomly selected on each trial.

**Figure 1.**
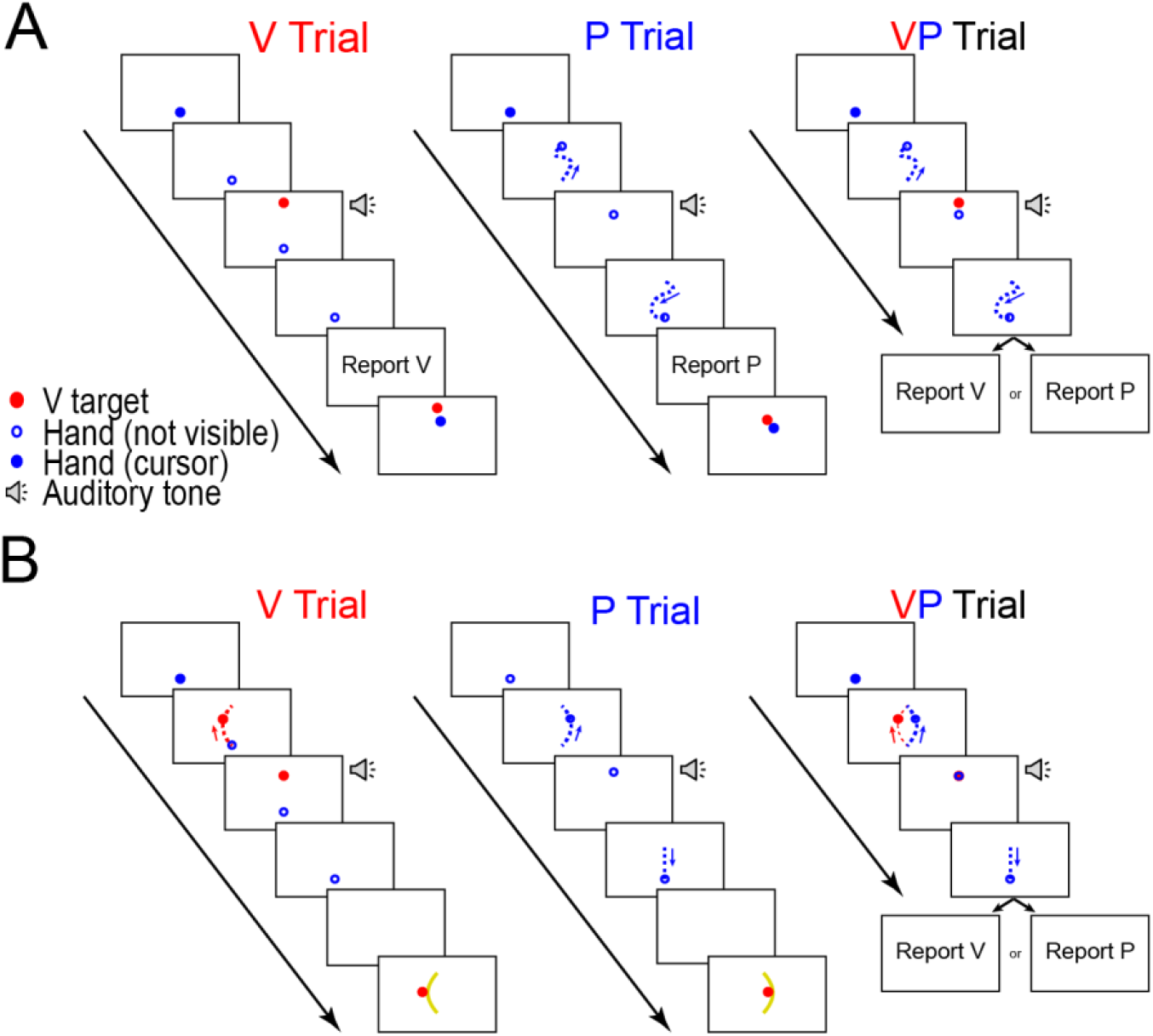
Schematic of the paradigm for Experiment 1. (A) The static estimation task had participants reproduce a stationary position in the task space. The target position was either cued visually with a circle appearing on the display (V Trial), or it was cued proprioceptively as the position of the index fingertip at the endpoint of a random passive movement (P Trial). The task began with a baseline block of unimodal (V and P) trials, followed by a block of both unimodal and bimodal (VP) trials. In all trials, the visual target was located 6 cm further from the participant than the proprioceptive target. In VP trials, participants were presented with both visual and proprioceptive cues. They were informed of the offset between cues and were instructed (prior to the trial) as to which sensory modality they should attend and report. Report accuracy feedback was provided only on unimodal V and P trials. (B) The dynamic estimation task had participants reproduce a curved movement trajectory in the task space. The trajectory was cued either visually with the movement of a circular cursor (V Trial), or it was cued proprioceptively as the path traveled by the index fingertip during passive movement of the arm (P Trial). The task began with a baseline block of unimodal V and P trials, which was followed by a test block that included both unimodal and bimodal (VP) trials. In all trials, the visual and proprioceptive trajectories comprised opposing curves offset by 6 cm at the midpoint. As in the static estimation task, participants were informed of the offset and were instructed prior to VP trials as to which sensory modality they should attend and report. Midpoint accuracy feedback was provided only on unimodal trials.

### Static Estimation Task

Figure 1a shows a schematic of the static estimation task. Here, participants experienced 4 types of trials modeled after tasks by Block and colleagues (Block and Bastian, 2011, 2010). The first 2 trial types tested unimodal sensory estimation. In Visual (V) trials, participants were shown a stationary visual target 18 cm in front of the starting position and heard an auditory tone cueing them to focus on the target location. After 1000 ms, the target vanished, and participants were asked to move their dominant hand to align the index finger to the remembered target location. In Proprioceptive (P) trials, participants’ dominant hands were passively moved to align the index fingertip with a stationary endpoint 12 cm in front of the starting position (6 cm away from the V target) via a randomly selected Bezier curved path constructed from 4 control points. They were then presented with an auditory tone cueing them to focus on the target position. After 1000 ms had elapsed, the hand was returned to the starting position along another random Bezier curve. The random Bezier curves prevented participants from using information about the passive movement to supplement the estimation of the target endpoint position. Once the hand was passively returned to the start position, participants were allowed to move their hand to re- align their index fingertip to the previously perceived endpoint. In the unimodal V trials and P trials, participants received visual feedback about the accuracy of their endpoint localization relative to the actual target position to stabilize perception in these trials and to reduce the degree of sensorimotor recalibration that may have occurred during the task. No visual feedback was provided during hand movements.

The remaining 2 trial types tested bimodal (i.e., visual and proprioceptive) sensory estimation. At the start of these trials, participants were instructed to attend to one sensory modality and ignore the other. The unseen dominant hand was passively moved along a random Bezier curve to align the index fingertip with a stationary endpoint. Once at the endpoint, the visual target was also displayed, and an auditory tone cued participants to focus. After 1000 ms, the visual target was extinguished, and the hand was returned to the starting position along another random Bezier curve. Participants were then instructed to move their unseen hand to align the index fingertip with the visual target (VPV trials) or the proprioceptive endpoint (VPP trials), depending on which sensory modality had been cued at the trial’s start. No feedback of reporting accuracy was provided here. As participants were explicitly instructed on which sensory modality they should focus and report, systematic errors in the direction of the “ignored” target could be attributed to unintentional biases induced by the other modality.

The visual target and proprioceptive endpoint were misaligned in all trials such that the visual target was 6 cm further from the subject than the proprioceptive endpoint, and participants were informed of the offset prior to beginning the task. This separation was chosen to be similar to prior studies (Block and Bastian, 2011, 2010) and to be robust relative to typically observed midpoint and endpoint variability across individuals without being so large that people would be unlikely to integrate the two positions into a single percept (Hsiao et al., 2022). In all cases, individuals were required to remember the cued location rather than report the location of an existing target, which enabled us to match the memory requirements of the static estimation task to those of the dynamic estimation task described below.

The static estimation task consisted of two blocks of trials. In the first block (120 trials), participants experienced an equal number of unimodal V trials and unimodal P trials in a random order. The first block served as a baseline and allowed us to estimate perceptual reports for the unimodal trials alone. The second block comprised a mix of unimodal (24 V and 24 P trials) and bimodal trials (60 VPV and 60 VPP trials), randomly intermixed.

### Dynamic Estimation Task

Figure 1b shows a schematic of the dynamic estimation task. Here, all outward movements of the visual stimulus and the hand occurred along symmetric arc-shaped Bezier curves with 3 control points, with all leftward-curved arcs associated with cues in one sensory modality and all rightward-curved arcs associated with cues in the other sensory modality. The pairing of arc direction and sensory modality was counterbalanced across participants. Half of the participants experienced visual cues on the left and proprioceptive cues on the right, while the other half experienced the opposite. The midpoint of each arc was displaced 3 cm from the midline, corresponding to a total horizontal displacement of 6 cm between the two arc midpoints. Each arc trajectory ended at the same point, 15 cm away from the start position. Movement along the arc was completed in 1600 ms (analogous to the timing of the static estimation task). Hand and cursor velocity both followed a bell-shaped velocity profile, resulting in a peak velocity of 0.187 m/s at the movement midpoint.

As in the static task, the dynamic estimation task comprised four trial types. The first two trial types tested unimodal sensory estimation. In V trials, participants watched a visual cursor move along an arcing trajectory. At the end of the arc, the cursor paused for 500 ms before being extinguished. Participants were then presented with an auditory tone, which cued them to move their dominant hand to trace the observed motion with their index fingertip. In P trials, the dominant hand was passively moved such that the index fingertip traced an unseen arcing trajectory and was held at the endpoint for 500 ms. The hand was then passively returned to the starting position in a straight-line path. Participants were instructed to move the unseen hand to reproduce the perceived trajectory of the index fingertip. Feedback was provided at the end of unimodal trials by displaying the target arc, along with a cursor showing the reproduced midpoint. The dynamic estimation task also tested two bimodal trial types, in which participants were instructed to attend to and report either the visual (VPV trials) or the proprioceptive (VPP trials) trajectory and ignore information from the other sensory modality. The timing of events during each trial, as well as passive and active movement speed ranges, were matched to those of the static estimation task. Any observed differences between the static and dynamic estimation tasks were, thus, unlikely to result from differences in working memory requirements or speed of movement.

The dynamic estimation task consisted of two blocks. In the first baseline block (60 trials), participants experienced only unimodal V and P trials in random order. The second block consisted of a mix of unimodal and bimodal trials (20 V, 20 P, 60 VPV, and 60 VPP trials) presented in random order.

Prior to the start of block 1, participants were exposed to a series of familiarization blocks to ensure they could reasonably reproduce the arc trajectory and stop near the desired endpoint without online visual feedback of their hand position. Participants first practiced making point-to-point reaches to a desired endpoint, starting with online cursor feedback of the hand and then practicing without that feedback (but with endpoint feedback to see their movement errors). They then learned to produce arcing motions that passed through an indicated midpoint target and stopped at an endpoint target; as before, they did this with and without online cursor feedback. Participants only began the main task once they were reasonably comfortable and accurate at producing the desired movement.

#### Experiment 2

Participants were asked to complete a single session in which they performed a static estimation task similar to Experiment 1. The primary difference between the static estimation tasks from Experiment 1 and Experiment 2 is that both the V target and P endpoint were now presented 15 cm in front of the start position and were displaced along the horizontal axis 3 cm to the left and right of the participant’s midline (for a total displacement of 6 cm between them). This offset matched the direction of the visual-proprioceptive discrepancy in the dynamic estimation task of Experiment 1. The pairing of target side (left or right) and sensory modality (vision or proprioception) was counterbalanced across participants. The timing within and across trials, the block structure and number of trials, and participant instructions were otherwise identical to the static estimation task of Experiment 1.

#### Experiment 3

Participants were asked to complete a single session in which they performed a dynamic estimation task similar to Experiment 1. In Experiment 1, the V and P stimuli comprised arc trajectories with a midpoint displaced 3 cm on either side of the participant’s midline. In Experiment 3, both the V and P trajectories curved to the right of the midline. For nine of the fifteen participants, the midpoint of the arc was displaced 3 cm rightward from the midline (close) on V trials, and 9 cm rightward (far) on P trials. The V and P displacements were switched for the other six participants. Individuals were randomly assigned to one of the two displacement conditions with equal likelihood, although these groups were not rebalanced after excluding individuals for failing to follow directions. All other aspects of the task remained identical to the dynamic estimation task of Experiment 1.

### Data Analysis

Reaches were analyzed offline using programs written in MATLAB (The MathWorks, Natick, MA, USA). Data, analysis code, and model code are available at https://osf.io/wt6hz (DOI: 10.17605/OSF.IO/WT6HZ). For trials in both static estimation tasks, the movement endpoint was determined as the hand position along the x- and y-axis after the hand velocity remained below 0.06 m/s for 500 ms (selected to identify the first time the hand came to a stop and exclude any subsequent motor corrections). For trials in the dynamic estimation task, the movement midpoint was defined as the horizontal position of the hand when the hand had moved halfway (i.e., 7.5 cm along the y-axis) from the starting position to the ideal endpoint. The movement endpoint was determined as the position after the x- and y-hand velocity remained below 0.05 m/s for 2000 ms, to allow time for the hand to come to a complete stop along both axes (note, the movement midpoint is the position of interest in this task). For both the static and dynamic estimation tasks, the movement endpoint (y-axis position when the hand was stationary in Experiment 1, or x-axis position for Experiment 2) and midpoint (x-axis position when the y-axis position of the hand was 7.5 cm in Experiments 1 and 3), respectively, was used to calculate a report error, defined as the difference between the actual reported position and the ideal position. For unimodal trials, the report error was calculated separately for the baseline (block 1) and mixed trial (block 2) blocks. In bimodal trials, report error was calculated using the target/trajectory modality participants were instructed to reproduce. Positive report errors in bimodal trials reflected a reported position that was shifted toward the cued position of the other modality. Biases on bimodal trials (i.e., the extent to which the hand position was shifted by the presence of discrepant feedback from another sensory modality) were calculated as the difference between the reported position on those trials and the average reported position in corresponding baseline unimodal trials (e.g., VPV – Vaverage, baseline). Positive biases reflect a shift toward the ideal location of the other modality. For all tasks, movement peak velocity was identified as the greatest hand-speed vector magnitude achieved during the movement.

Differences in hand position (endpoint, midpoint) or biases (VPV – V and VPP – P) across trial types, both within individual tasks as well as across tasks, were analyzed with mixed-effects regression models using the lme4 package in R (Bates et al., 2015). Models included a random intercept of subject and a random slope of trial type by subject. Factors were tested for significance using likelihood ratio tests comparing models with and without the factor of interest. Posthoc comparisons were examined using the emmeans package in R with Kenwood-Roger estimates of degrees of freedom and corrected for multiple comparisons using the Tukey method (Lenth et al., 2021). Models that included the dynamic estimation task for Experiment 1 initially included a nuisance factor of Side (V-left or V-right), which was removed from the final analysis as it showed no significant effects (*p* > 0.59). Side was found to be significant in models for Experiment 2 (p = 0.02), likely due to biomechanical factors, so this factor was left in the model for those analyses. Side (V-close or V-far) showed no significant effect for Experiment 3 and was removed from the final analysis (*p* = 0.35). Models in Experiment 1 that included both static and dynamic estimation tasks initially also included a nuisance factor of task order, which was removed from the final analysis because it showed no significant effects (*p* > 0.28). Although individuals were asked to make their report movements within a desired speed range, all models included a nuisance factor of peak velocity to control for differences in movement speed. The peak velocity of report movements was found to be significant in Experiment 1 (*p* < 0.001) and Experiment 3 (p < 0.001), but was not significant for Experiment 2 (*p* = 0.71). Hence, peak velocity was retained in the regression models for Experiments 1 and 3, but not Experiment 2. Finally, in some cases, simple regressions were performed to examine the relationships among variables of interest and/or for data visualization purposes.

To confirm the validity of our multisensory integration model, we calculated an estimate of each individual’s ideal bimodal variance based on the variances of vision and proprioception observed during baseline unimodal trials according to equation (3). We then tested the relationship between the estimated bimodal variance and the largest mean bias exhibited in bimodal trials of the static and dynamic estimation tasks (regardless of sensory modality) from Experiment 1, under the hypothesis that larger biases should reflect a greater tendency for that individual to rely on multisensory integration.

## RESULTS

### Modeling

When computing a state estimate based on sensory feedback, it is generally assumed that individuals rely on multisensory integration. When the limb is stationary, multisensory integration presents no concerns; since the limb position remains constant, feedback delays are of little consequence. Hence, multisensory integration will provide an accurate estimate of the current body state in this case. The most widely accepted model of multisensory integration, the maximum likelihood estimation model (Ernst and Banks, 2002; van Beers et al., 1996), explains this benefit clearly. When the limb is moving, however, sensory delays mean that any sensory feedback is now out of date, and relying on it will increase the uncertainty of a state estimate. Passive movement of the limb is a particularly interesting situation to consider here, as there is no motor command efference copy from which the motor system can make predictions about the current body position (Miall and Wolpert, 1996). In such cases, we hypothesized that it may be more beneficial to rely on unimodal feedback to avoid uncertainty arising from differences in sensory delays across modalities.

To first visualize the effect of sensory delays on the reliability of multisensory integration, we updated to the maximum likelihood model (see Methods and Fig. 2). Rather than assuming one knows the current limb position at each instant in time, we instead assume that one only knows the limb position sometime in the past, with each sensory modality potentially reporting a different limb position based on differences in sensory delays. To simulate this, we included terms in the model that reflected the perceived limb position sometime in the past, where the difference in limb position was calculated according to the sensory delay and the current limb speed (see Methods, Equation 1). Note that we did not make any changes to the sensory uncertainty terms of the original model: we assumed that the perceived limb position did not degrade over time, and that sensory uncertainty did not scale with movement speed. We also made no assumptions regarding the magnitude of any intrinsic visual or proprioceptive biases in limb position.

**Figure 2.**
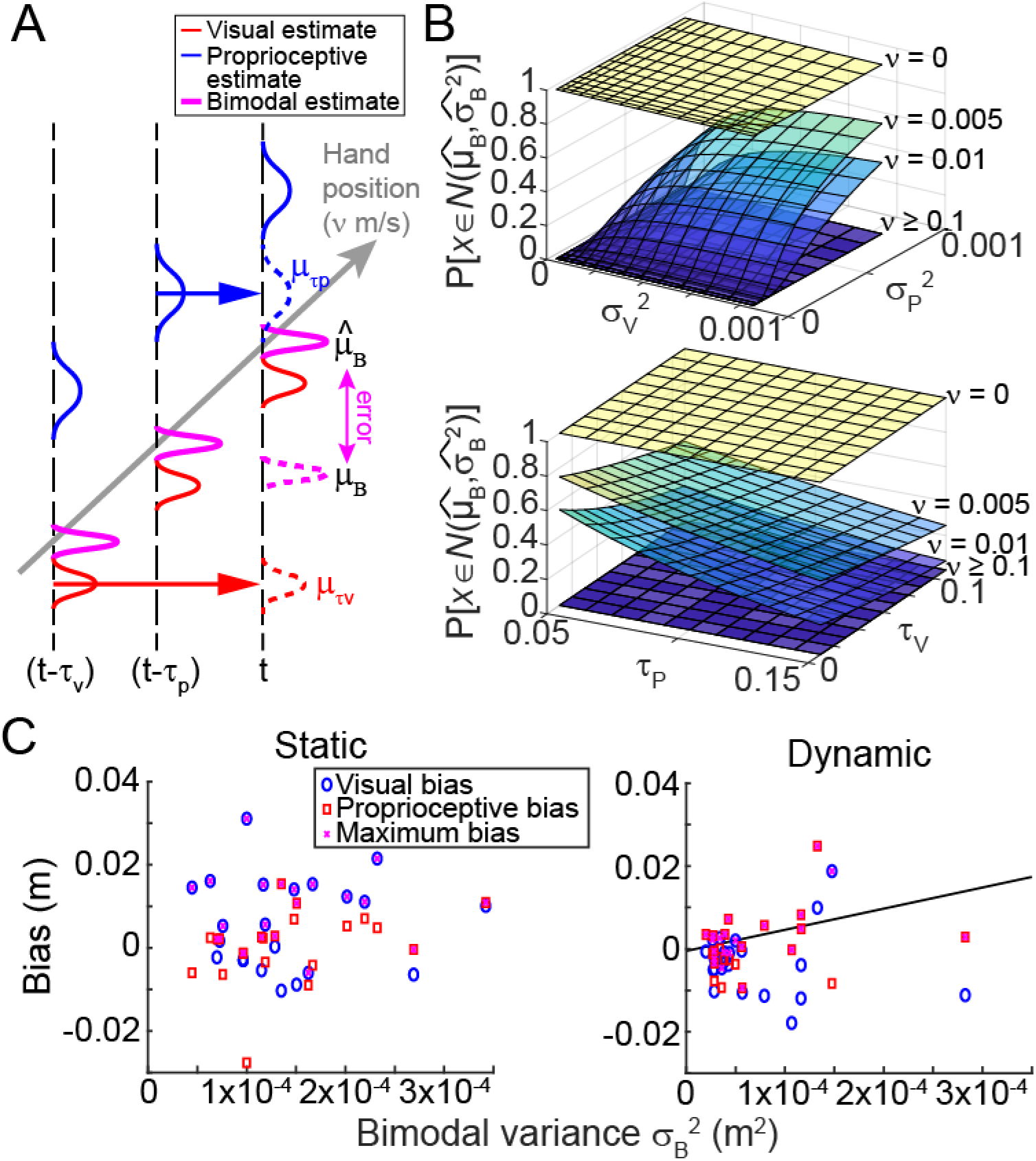
Model results. (A) Updated MLE model accounting for limb movement and sensory delays. If the limb moves with speed, ν, at time, t, we have delayed visual (μτv, red dashed line) and proprioceptive (μτp, blue dashed line) feedback of the limb from sometime in the past instead of at its current position (red and blue solid lines). This leads to an erroneous bimodal state estimate (μB, purple dashed line) that is offset from the ideal bimodal estimate (𝜇^𝐵, purple solid line) by some error (purple arrow). (B) Simulation results suggest that as movement speed (ν) increases, the probability of obtaining an accurate bimodal sensory estimate decreases; in such cases, it may be better to use a unimodal state estimate. However, this also depends on the relative uncertainties of the two sensory modalities (top panel) and the relative time delay between the two modalities (bottom panel). (C) Observed data from Experiment 1 was used to estimate the bimodal variance via the MLE combination of the observed unimodal sensory variances. For the static estimation task (left), no relationship was observed between the bimodal variance and the observed bias. In contrast, in the dynamic estimation task (right) there was a clear correlation across individuals. Consistent with the model, this suggests that individuals with greater bimodal variance in the dynamic estimation task likely relied on an integrated state estimate and were more susceptible to discrepant information from the other sensory modality.

Model simulations showed that when estimating the position of a stationary limb, bimodal integration offered the greatest reliability, replicating previous work (Berniker and Kording, 2011; Ernst and Banks, 2002; van Beers et al., 1999, 1996). Our simplistic model suggested that the advantage afforded by bimodal state estimation persisted at very slow movement speeds (< 0.01 m/s), particularly when the uncertainty of the individual sensory modalities was high or when sensory delays were short. As movement speed increased, however, the bimodal sensory estimate became quite poor (i.e., there was no overlap between the Gaussian representing the estimate given sensory delays and that of the ideal estimate assuming the system had access to the current limb position). The error arose because the state estimate was reliant on temporally delayed sensory feedback informing the system of where the limb was located some time ago rather than its current location. Under these circumstances, the model suggests that it may be better to use unimodal sensory feedback rather than integrated multisensory feedback to derive a state estimate (Fig. 2B). This prediction is in line with those arising from more complex models that also consider active (i.e., voluntary) movement of the limb (Crevecoeur et al., 2016; Kasuga et al., 2022; Oostwoud Wijdenes and Medendorp, 2017). To test this key prediction of our model, we examined how individuals estimated their current body state while stationary versus when being passively moved.

#### Experiment 1

This experiment examined the degree to which individuals could report limb state using sensory information from one modality while ignoring discrepant information from another. Participants completed four trial types. In the two unimodal trial types, participants were required to report the perceived location of a visual or a proprioceptive target with their hidden dominant hand (V and P trials, respectively). In the two bimodal trial types, the visual and proprioceptive targets were presented together. Here, participants were explicitly cued to report only the visual target and ignore proprioceptive information (VPV trials), or report the proprioceptive target and ignore vision (VPP). Performance was measured by examining the accuracy with which the perceived movement endpoint (static task) or midpoint (dynamic task) was reported. If individuals relied on multisensory integration to inform their state estimate, we expected to observe positive report errors on bimodal trials relative to baseline unimodal trials, reflecting a shift of perceived position toward the discrepant sensory modality. We likewise expected to observe positive report biases, computed as the difference between the report error on each bimodal trial relative to the average report error on corresponding baseline unimodal trials. Importantly, our model predictions suggested that we should observe more positive report errors and biases in the static estimation task compared to the dynamic estimation task.

### Limb state estimates at movement endpoint were biased by multisensory integration despite explicit instruction to ignore one sensory modality

In the static estimation task (Fig 1A), static visual cues were displaced 6 cm farther along the y-axis compared to proprioceptive movement endpoints (Block and Bastian, 2011, 2010). Individuals were asked to report the perceived locations of these cues. Report error was significantly different across the four types of trials (Fig. 3A,B; χ^2^(3) = 10.51, *p* = 0.015). This effect was driven by a difference between VPV and V trials (*p* = 0.010), in which VPV trials exhibited a larger positive report error (i.e., they were shifted toward the location of the P cue). There was also a difference between VPV and VPP trials (*p* = 0.012), with VPV trials exhibiting more positive report errors than VPP trials. The difference between VPP and P trials (*p* = 0.54) and the difference between V and P trials (*p* = 0.058) did not reach significance. At the individual subject level (Fig 4A), there was no relationship observed between the bias in the Visual modality and the bias in the Proprioceptive modality (*r*^2^ = 0.10, *p* = 0.13). Together, these results suggest that when reporting the location of visually cued targets, individuals have difficulty ignoring discrepant proprioceptive feedback when the limb is stationary.

**Figure 3.**
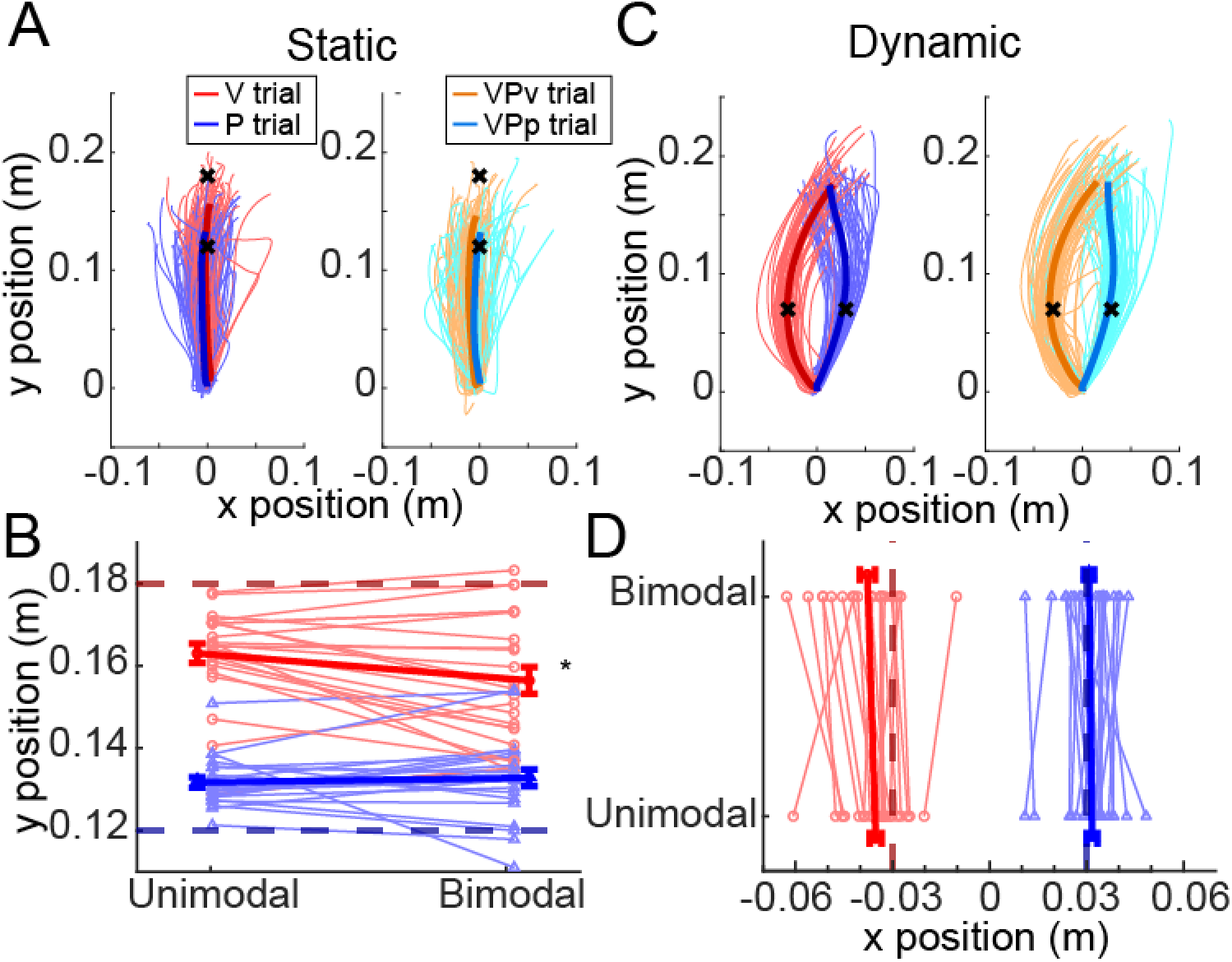
Results from the static and dynamic estimation tasks of Experiment 1. (A) Data from a single participant in the static estimation task performing baseline unimodal trials (left panel) and bimodal trials (right panel). The cued endpoint location is indicated by a black x. (B) The average reported endpoint position for each participant is shown along with the group mean for visual (V and VPV, red) and proprioceptive (P and VPP, blue) trials. (C) Data from the same participant depicted in panel A performing unimodal (left) and bimodal (right) trials of the dynamic estimation task. (D) Group data for the Dynamic task akin to panel B. All units are in meters.

**Figure 4.**
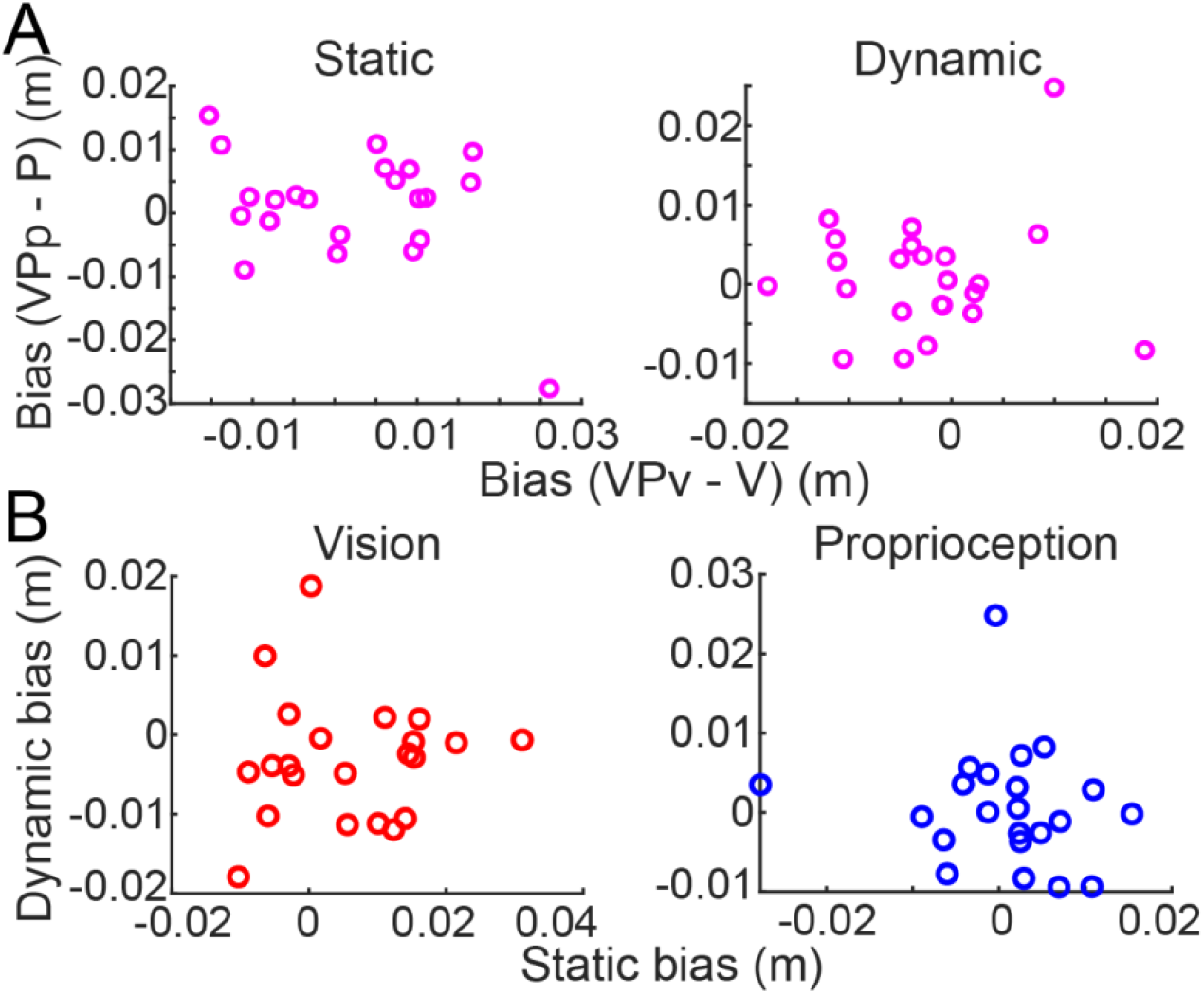
Report biases in Experiment 1. (A) Across individuals, there was no relationship between the biases observed when reporting visual and proprioceptive cues in either the static (left panel) or dynamic (right panel) estimation tasks. (B) Across the static and dynamic estimation tasks, there were no relationships between the biases observed on bimodal trials in response to the visual cue or in response to the proprioceptive cue.

To determine whether the difference in report errors between VPV and VPP trials was related to the baseline variance displayed in the visual and proprioceptive modalities, we assessed the relationship between the report error variance exhibited in unimodal trials and the bias in bimodal trials. Within the visual modality, there was no relationship between the variance in unimodal V trials and bias in VPV trials (*r*^2^ = 0.04, *p* = 0.37). Similarly, there was no relationship between the variance in unimodal P trials and bias in VPP trials (*r*^2^ = 0.05, *p* = 0.30). Peak velocity was also consistent across all trial types (χ^2^(1) = 0.02, *p* = 0.88). These results suggest that, overall, the observed difference between VPV and VPP trials was not due to baseline variance in either modality or a difference in the kinematics of report movements.

### Limb state estimates were not biased by multisensory integration during movement

For the dynamic estimation task (Fig 1B), individuals experienced arcing motion patterns that curved either to the right or left, with visual cues consistently arcing in one direction (e.g., rightward) and proprioceptive cues arcing in the other direction (e.g., leftward). Arc midpoints were separated by 6 cm. Individuals were asked to report the perceived motion on each trial, with a focus on reproducing the midpoint of the movement. Across all participants, there was no significant effect of the pairing of arc direction and sensory modality on report error (χ^2^(1) = 0.26, *p* = 0.61), so data were analyzed together (data were collapsed across participants who experienced V trials that curved rightward and those who experienced the V trials that curved leftward).

In contrast to the static estimation task, report error (Fig 3C,D) here showed no significant differences across the four trial types (χ^2^(3) = 4.49, *p* = 0.21). There was no relationship between the bias observed in the Visual modality and the bias observed in the Proprioceptive modality (Fig 4A; *r*^2^ = 0.01, *p* = 0.69). There was also no relationship between the variance in report error exhibited in baseline unimodal trials and the bias observed in bimodal trials in either sensory modality (Vision: *r*^2^ = 0.02, *p* = 0.48; Proprioception: *r*^2^ = 0.08, *p* = 0.18). Together, these results suggested that individuals were not susceptible to the influence of discrepant sensory feedback when reporting limb movement.

There was no systematic difference in the peak velocity of report movements across the different trial types (interaction: χ^2^(3) = 0.35, *p* = 0.94). However, there was a significant negative effect of peak velocity on report error, in which the report error became more positive (i.e., the movement curvature decreased) as peak velocity decreased (χ^2^(1) = 172.46, *p* < 0.001).

### Static state estimation was significantly different and independent of dynamic state estimation

Comparing report errors in the static and dynamic estimation tasks, we first ruled out an effect of task order (χ^2^(1) = 1.16, *p* = 0.28). There was a significant negative effect of the peak velocity of report movements (χ^2^(1) = 153.01, *p* < 0.001) on report error, however, so this variable was included as a nuisance parameter in our regression model. The analysis showed a significant main effect of task (χ^2^(5) = 3235.8, *p* < 0.001) wherein participants showed more positive report errors (i.e., greater bias) in the static compared to the dynamic estimation task. There was also a significant main effect of trial type (χ^2^(7) = 424.08, *p* < 0.001), in which the report error was different for each trial type (post hoc comparisons all showed *p* < 0.001).

Finally, there was a significant interaction between task and trial type (χ^2^(3) = 414.68, *p* < 0.001), which was driven by differences in report error across trial types in the static estimation task that were not present in the dynamic estimation task. That is, for the static task, report error was more positive (i.e., biased more toward the direction of the other sensory modality) in V trials compared to P trials (*p* = 0.02), in VPV trials compared to V trials (*p* = 0.0001), and in VPV trials compared to VPP trials (*p* = 0.0004). In contrast, for the dynamic task, report error was significantly less positive in VPV trials compared to VPP trials (*p* = 0.03), with no observed differences between the other trial types (*p* > 0.12). These findings align with the within-task results those reported above.

There was no relationship between biases shown in bimodal trials of the static and dynamic estimation tasks for either the visual or the proprioceptive modality (Visual bias: *r*^2^ = 0.00, *p* = 0.83; Proprioceptive bias: *r*^2^ = 0.02, *p* = 0.49, Fig. 4B). Additionally, the variance that participants exhibited on unimodal trials at baseline was not correlated across tasks (i.e., individuals were not consistently variable across tasks), for either vision or proprioception (Visual variance: *r*^2^ = 0.15, *p* = 0.07; Proprioceptive variance: *r*^2^ = 0.06, *p* = 0.25). Thus, individual performance in the static and dynamic estimation tasks was uncorrelated, consistent with the analysis of report error above.

### The visual-proprioceptive conflict in bimodal trials recalibrated unimodal perception in the static but not the dynamic estimation task

Since participants received no performance feedback on bimodal trials, unimodal trials with feedback were presented in block 2 to keep participants’ movements relatively stable. The presence of these unimodal trials afforded an opportunity to examine whether the sensory conflicts in bimodal trials would also modulate the performance of unimodal trials (e.g., through recalibration; Fig 5). Within the static estimation task, we observed a significant main effect of block (χ^2^(2) = 12.90, *p* = 0.002), with unimodal trials in block 2 being more positively biased compared to the baseline block 1. There was no effect of trial type (V versus P trials; χ^2^(2) = 3.40, *p* = 0.18) or block-by-trial-type interaction (χ^2^(1) = 0.12, *p* = 0.73), suggesting that the biasing of unimodal trials was similar across both sensory modalities. In contrast, for the dynamic estimation task, we observed no effect of block (χ^2^(2) = 0.94, *p* = 0.62) or trial type (χ^2^(2) = 1.70, *p* = 0.43), nor was there an interaction (χ^2^(1) = 0.003, *p* = 0.95). These findings suggest that exposure to conflicting sensory cues induced biases toward the opposing sensory modality for both bimodal and unimodal trials in the static but not the dynamic estimation task. Notably, these biases were observed even though conflicting sensory cues were not present during unimodal trials. This may be an indication that sensory recalibration occurred in the static but not the dynamic estimation task.

**Figure 5.**
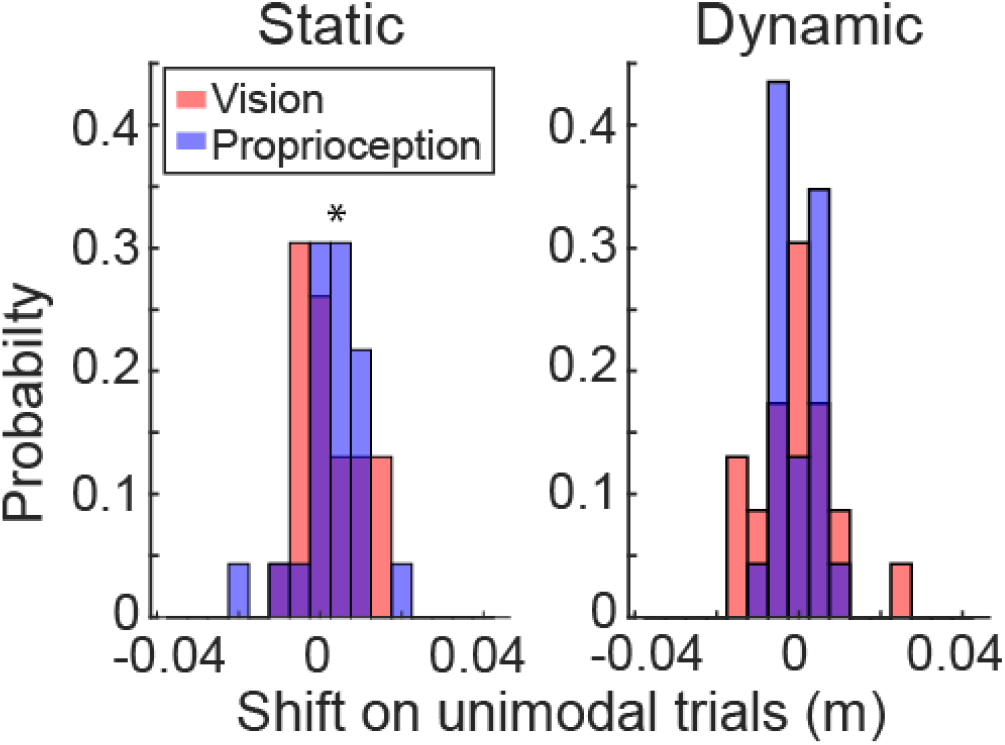
Shift in report error for unimodal trials in Experiment 1. In the static estimation task (left panel), participants exhibited a shift in report error on unimodal trials in the context of conflicting bimodal sensory cues relative to baseline. In contrast, unimodal trials in the dynamic estimation task exhibited no such shift when bimodal trials were introduced.

### The bias displayed in bimodal trials correlated with overall sensory uncertainty in the dynamic but not the static estimation task

Although we found that the bias shown in bimodal trials was not correlated with the variance of unimodal sensory estimates at baseline, our computational model predicted that the bias shown in bimodal trials should correlate with the magnitude of overall sensory uncertainty. Importantly, this relationship was predicted to be idiosyncratic across state estimation contexts. Greater sensory variance overall widens the Gaussian representing the bimodal state estimate (i.e., the calculated variance of the bimodal estimate would be larger, see Methods, Equations (3) and (4)), increasing the likelihood that any bimodal estimate would fall within the bounds of the ideal estimate had current (i.e., non-delayed) sensory information been available. As temporal delays are problematic for the dynamic, but not the static state estimation context, the model predicted a positive relationship between bias in bimodal trials and overall sensory variance for the dynamic context alone.

To check this prediction, we examined the biases exhibited by individual participants in both the static and dynamic estimation tasks (Fig. 2C). Although people were only significantly biased in the static estimation task, there was variability across individuals, with some showing a bias in the dynamic task as well. We compared the largest bias individuals exhibited in the dynamic task, regardless of sensory modality, against the estimated bimodal variance calculated using the variances exhibited in baseline V and P trials (see Methods, equation (3)). Consistent with the model prediction, we found a positive correlation between the theoretical bimodal variance and the extent to which individuals exhibited a bias in the dynamic estimation task (*r*^2^ = 0.18, *p* = 0.047; after removing the one outlier participant: *r*^2^ = 0.47, *p* = 0.001). When an individual’s ideal bimodal variance was quite large, their reports of limb movement were more strongly biased by the discrepant sensory modality they were asked to ignore. On the other hand, individuals with low bimodal variance tended to exhibit less bias in the dynamic estimation task. In contrast, during the static estimation task, individuals all exhibited a bias regardless of their sensory uncertainty (*r*^2^ = 0.004, *p* = 0.78), suggesting that all participants in the static estimation task favored a bimodal state estimate regardless of their bimodal variance because their movement speed was zero.

#### Experiment 2

While the static estimation task in Experiment 1 was based on tasks used in prior work (Block and Bastian, 2011, 2010; Reuschel et al., 2010), it differed from the dynamic estimation task in two ways. First, it involved reporting stationary rather than dynamic positions of the limb, the key manipulation in that study. Second, the static and dynamic estimation tasks presented the dissociation between the visual and proprioceptive cue locations in different directions relative to the primary direction of limb movement. In the static estimation task, cues from the two sensory modalities were displaced along the primary direction of limb movement. In contrast, the dynamic estimation task created a discrepancy between the two sensory modalities in a direction orthogonal to the primary movement direction. To determine whether this latter difference drove the results of Experiment 1, we asked a new group of individuals to complete a version of the static estimation task in which the cued visual and proprioceptive locations were displaced in the horizontal direction – orthogonal to the primary movement direction of the hand and analogous to the displacement introduced during the dynamic estimation task. As in Experiment 1, we expected to see positive report errors and biases toward the direction of the discrepant sensory modality if individuals are relying on bimodal state estimates.

### Static state estimation was biased by a visual-proprioceptive conflict in the direction orthogonal to the axis of movement

When analyzing report error in Experiment 2, we observed a significant effect of testing side (χ^2^(1) = 5.12, *p* = 0.02), likely due to rightward movements primarily involving elbow movement while leftward movements involved both elbow and shoulder movement. Testing side was retained in the regression model to account for this difference. There was no effect of peak velocity (χ^2^(1) = 0.14, *p* = 0.71). Critically, there was a significant difference in report error across the four trial types when comparing unimodal trials at baseline to bimodal trials (χ^2^(3) = 12.21, *p* = 0.007, Fig 6A). The effect was driven by a difference between VPV and V trials (*p* = 0.016) and a difference between VPP and P trials (*p* = 0.039). In both cases, we observed a larger positive report error in bimodal trials, suggesting that report errors were shifted toward the location of the discrepant sensory modality. There was no significant difference observed between VPV and VPP trials (*p* = 0.54), or between baseline V and P trials (*p* = 0.20). We also observed no difference in unimodal trials before and after the introduction of the bimodal trials (*p* > 0.17). At the individual subject level, there was no relationship observed between the bias in the visual modality and the bias in the proprioceptive modality (*r*^2^ = 0.0001, *p* = 0.97). Within sensory modality, there was no relationship observed between the report error variance exhibited in baseline unimodal trials versus the bias in bimodal trials (Vision: *r*^2^ = 0.01, *p* = 0.68; Proprioception: *r*^2^ = 0.0003, *p* = 0.94). These results suggest that when reporting the location of static targets, individuals have difficulty ignoring discrepant feedback from the other sensory modality, even when the sensory cues are displaced orthogonal to the primary direction of movement.

**Figure 6.**
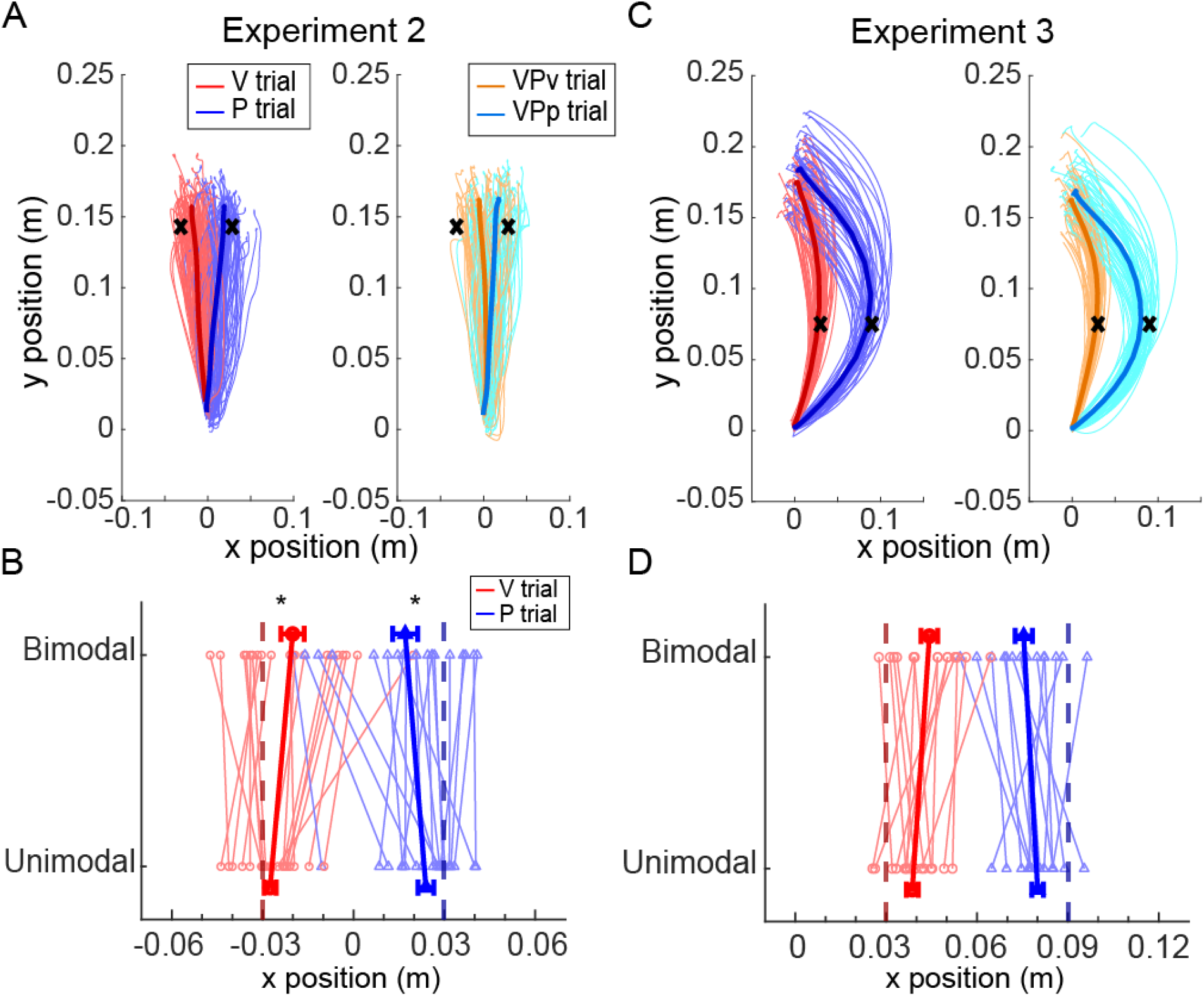
Report errors in Experiments 2 and 3. (A) In Experiment 2, participants completed a static estimation task in which the sensory cues were displaced orthogonal to the primary direction of movement. Data from a single subject performing unimodal V and P trials (left panel), and bimodal VPV and VPP trials (right panel) are shown. Thick lines reflect the average movement for each condition across trials. The cued endpoint location is indicated by a black x. (B) The average reported endpoint positions for each participant in Experiment 2 (thin lines) are shown along with the group mean (thick lines), for visual (V and VPV, red) and proprioceptive (P and VPP, blue) trials. Overall, participants exhibited a bias in report error on bimodal trials relative to unimodal trials when reporting the location of both visual and proprioceptive cues. (C) In Experiment 3, participants completed dynamic estimation task in which the sensory cues were displaced on the same side of the midline, causing movement paths to be curved in the same direction. Single-subject data are shown for unimodal (left panel) and bimodal trials (right panel). The cued midpoint location is indicated by a black x. (D) Average reported midpoints for each participant (thin lines) are shown along with the group mean (thick lines). Participants did not exhibit any significant bias in report error on bimodal trials relative to unimodal trials when responding to either the visual or proprioceptive cues. All units are in meters.

#### Experiment 3

The dynamic estimation task in Experiment 1 consisted of arcing movements with opposing curvatures for each of the sensory modalities, which may have facilitated the separation of visual and proprioceptive stimuli. To control for this possibility, we conducted a third experiment in which the V and P arc trajectories were both curved in the same direction (i.e., both curving to the right of a participant’s midline) but with different magnitudes of midpoint displacement from the midline. In line with our findings from Experiment 1, we expected to see no difference in report error for bimodal trials compared to baseline unimodal trials if people can successfully ignore discrepant feedback from an opposing sensory modality when forming a dynamic state estimate.

### Dynamic state estimation was not biased by a visual-proprioceptive discrepancy when movement direction was matched

Although we observed no significant effect of testing side (i.e., whether the V or the P sensory modality was displaced further from the midline, χ^2^(1) = 0.86, *p* = 0.35), there was a significant effect of the peak velocity of report movements (χ^2^(1) = 56.48, *p* < 0.001), and a significant interaction between peak velocity and testing side (χ^2^(3) = 35.48, *p* < 0.001). As stated in the Methods, peak velocity was retained as a factor in our regression analysis for this experiment. The interaction likely reflected the differing degrees of curvature for movements along the close versus far movement paths (Zago et al., 2018).

Our regression analysis showed no significant effect of trial type on report error (χ^2^(3) = 2.21, *p* = 0.53; Fig. 6B). This suggested that movements to report perceived visual or proprioceptive cues were similarly accurate, regardless of whether there was also a discrepant cue provided with the opposing sensory modality. We also observed no difference in report error between V and P trials during baseline (χ^2^(1) = 1.21, *p* = 0.27). When considering only the unimodal trials across the entire experiment, we observed an effect of block (χ^2^(1) = 9.83, *p* = 0.007), in which the bias observed on V and P trials shifted away from (rather than toward) the discrepant sensory modality following the introduction of bimodal trials (*p* = 0.004). There was no effect of trial type (χ^2^(1) = 3.11, *p* = 0.21) or interaction between trial type and block (χ^2^(1) = 2.19, *p* = 0.14) in unimodal trials. At the individual subject level, there was no relationship observed between the bias in the visual modality and the bias in the proprioceptive modality (*r*^2^ = 0.01, *p* = 0.79). Within sensory modality, there was no relationship observed between the report error variance exhibited in baseline unimodal trials and the bias in bimodal trials (Vision: *r*^2^ = 0.21, *p* = 0.09; Proprioception: *r*^2^ = 0.01, *p* = 0.67). These findings replicate the results of Experiment 1, suggesting that people are able to ignore discrepant information from an opposing sensory modality when moving. Importantly, it does not seem to matter whether a conflict between visual and proprioceptive cues arises from a displacement of those cues across the midline (causing a switch in the direction of the movement curvature) or a difference in the magnitude of displacement on the same side of the midline.

## DISCUSSION

Estimating the current limb state is critical for proper motor control and uses both feedforward and feedback information. Multisensory integration is thought to be a key process supporting feedback-based state estimation, but studies of the phenomenon have largely focused on estimating static limb position. Due to differing temporal delays across sensory modalities, multisensory integration may be less useful when estimating limb movement (i.e., a dynamic context in which the limb position is constantly changing) (Cluff et al., 2015; Crevecoeur et al., 2016; Kasuga et al., 2022; Oostwoud Wijdenes and Medendorp, 2017). In this series of experiments, we examined whether multisensory integration for state estimation is obligatory by asking to what extent feedback from one modality interfered with the attempt to estimate the state of the limb according to a different sensory modality.

### Discrepant sensory feedback informs body state estimation differently when stationary or moving

Our results show that it is difficult to avoid multisensory integration when estimating static limb position, even when individuals know that the information conveyed by one sensory modality is task-irrelevant. We found that the effect persisted regardless of whether the sensory discrepancy aligned with or was orthogonal to a movement that brought the limb to its stationary position, suggesting that factors like the direction of offset may not constitute a useful heuristic when gauging feedback salience. Across individuals, the biases observed when people were asked to report the visual cue and ignore proprioceptive information were unrelated to biases exhibited when instructed to do the opposite. Additionally, the degree to which individuals were biased by multisensory integration in bimodal trials was unrelated to their estimation variance in each modality at baseline. Although this latter result runs counter to traditional minimum variance integration, other work has also shown similar results (e.g., Block & Bastian, 2010, 2011). Thus, our findings are generally consistent with those previously reported in the literature and in line with the hypothesis that multisensory integration is a useful strategy for sensory estimation of limb position (Berniker and Kording, 2011; Ernst and Banks, 2002; van Beers et al., 1996). Notably, our results extend this notion by suggesting that when estimating static limb position, multisensory integration may be difficult to explicitly avoid.

In contrast to the static estimation tasks, individuals exhibited no biases when reporting perceived passive limb movement in bimodal trials of the dynamic estimation task. That is, participants were able to accurately report either the visually or proprioceptively cued motion while successfully ignoring conflicting information from the other sensory modality. Any small bias that individuals exhibited when reporting dynamic motion was unrelated to their bias when reporting static position for either the visually cued or the proprioceptively cued limb state. We again found that the direction of multisensory discrepancy did not explain our results by showing that individuals could successfully ignore conflicting information regardless of whether the visual and proprioceptive trajectories had similar or opposing curvatures. Rather, the distinction in results between the static and dynamic estimation tasks was likely related to whether the limb is stationary or moving.

Overall, our results suggest that when the limb is moving, people are better able to isolate incoming information from a single sensory modality to form an estimate of the perceived limb state. This is reasonable, given that sensory information from vision and proprioception are delayed to different extents (delay of ∼20 ms to S1 vs ∼70 ms to V1 in macaques; Bair et al., 2002; Nowak et al., 1995; Raiguel et al., 1989; Song and Francis, 2013), and hence are effectively providing information about the limb at different points in the past. An instantaneous bimodal state estimate of the moving limb is likely to be less accurate than a state estimate based only on a single sensory modality when the limb is moving quickly. Indeed, others have observed a greater reliance on proprioceptive rather than visual feedback when the limb is moving (Crevecoeur et al., 2016; Kasuga et al., 2022). Hence, it is reasonable that state estimation during movement may use sensory information in a different manner compared to when the limb state is not significantly changing over time.

In Experiment 1, the bias observed on bimodal trials when reporting static position and the lack of bias when reporting limb movement were echoed in the report errors observed for unimodal trials. Unimodal trials were presented by themselves in block 1 (baseline) and were interleaved with bimodal trials in block 2 when individuals experienced discrepant sensory feedback. In the static estimation task, participants shifted the reported position of unimodal targets in block 2 relative to baseline, mimicking the bias observed in bimodal trials. In contrast, report errors for unimodal trials in block 2 of the dynamic estimation task show no such shift.

Although we did not observe a similar shift in unimodal trials for the static estimation task in Experiment 2, we observed a shift in bias away from the direction of the discrepant sensory modality in the dynamic estimation task of Experiment 3. Taken together, these results again indicate a difference between the degree to which unimodal state estimates are affected by feedback from a different, discrepant sensory modality in the static and dynamic tasks.

Specifically, these findings suggest that the nature of the feedback information used to compute a state estimate and the degree to which individual sensory modalities are recalibrated in the presence of a sensory mismatch may, at least partly, depend on whether this estimate is made while the limb is moving or stationary.

### Differences in multisensory integration when stationary or moving are predicted by a state estimation model accounting for sensory delays

Our behavioral results were consistent with the predictions of the mathematical model that motivated our study design. Our model was a modification of the conventional maximum likelihood estimation model (Ernst and Banks, 2002; van Beers et al., 1996) that accounted for limb movement and sensory delays. Model simulations suggested that individuals should favor a bimodal state estimate when stationary or moving very slowly since feedback delays would not cause wildly out-of-date estimates under these conditions. Our finding that report errors were biased in the direction of discrepant sensory information in the two static estimation tasks but not in the two dynamic estimation tasks supported this hypothesis. Our model also predicted increasing reliance on multisensory integration as overall sensory uncertainty increases when moving. These predictions were supported by an analysis of our study data from Experiment 1, showing that individuals with greater bimodal variance in the dynamic estimation task (i.e., individuals who exhibited greater uncertainty of their bimodal sensory estimates) were more likely to exhibit a positive bias in report error, suggesting that they continued to rely on multisensory integration to a small extent even when moving. This correlation was not observed in the static estimation task when all individuals were predicted to rely on bimodal state estimates regardless of their estimate variance. It would be interesting for future studies to test the predicted dependence of multisensory integration on movement speed more rigorously by asking individuals to report limb movement at different movement speeds.

In our behavioral task and model, we consider state estimation only in the case when the limb is passively moving. That is, we are specifically focusing on times when state estimation is reliant on sensory feedback alone. This stands in contrast to other studies in which individuals were asked to report their perception following actively generated movements (Block and Bastian, 2011, 2010; Block and Liu, 2023), in which state estimates could potentially additionally benefit from sensory predictions based on an efference copy of the ongoing motor command (Miall and Wolpert, 1996). Since sensory predictions provide an estimate of where the limb is expected to be at the current time, they can help overcome delays that would otherwise lead to out-of-date position estimates based on sensory feedback alone (Shadmehr et al., 2010).

Previous work has offered a more complex dynamic Bayesian model with Kalman filtering that attempted to account for sensory delays in an active arm movement task (Crevecoeur et al., 2016; Kasuga et al., 2022). Notably, these authors also concluded that individuals are likely to base state estimates on unimodal (in their case, proprioceptive) feedback in the face of sensory delays. Nevertheless, the contribution of sensory predictions to state estimation when the limb is actively moving and how that affects the integration of feedback from multiple sensory modalities needs further study.

When individuals appear to rely on a unimodal state estimate, our model does not make any strong predictions regarding which modality they should use. We suspect that the choice of modality may typically be determined by the relative sensory delays between vision and proprioception (and their effects on estimated limb position at different movement speeds), as well as their respective reliabilities. However, our findings suggest that task demands (i.e., instructions to ignore a particular sensory modality) can also modulate the choice of which modality to use. An interesting prediction of our model is that the choice of sensory modality may additionally depend on the direction of the baseline bias in that modality and the degree to which it aligns with the direction of the movement. That is, moving toward the direction of a sensory bias for a particular modality should improve the delayed feedback estimate coming from that modality, while moving away from the direction of the sensory bias should exaggerate the error. This prediction would need to be tested in future studies.

### Consideration of prior findings

Our findings are at odds with a prior study that showed individuals optimally integrate multisensory information when reporting the geometry of a perceived passive movement trajectory as following either an acute or obtuse angle (Reuschel et al., 2010). Specifically, the prior work demonstrated improved accuracy when reporting the angle under bimodal conditions as compared to only having vision or proprioception alone. This may seem to contradict our current findings; however, it is not clear whether individuals in the prior study relied on sensory integration throughout the entire movement. It is possible that in the prior task, individuals sensed the angle according to the positions of the movement endpoints or when the hand slowed down at the via point of the trajectory (i.e., when the hand changed direction to define the angle). In contrast, in our task, individuals were asked to reproduce the position at the midpoint of the passive movement, which occurred at the peak velocity when sensory delays were likely to have a large effect on the accuracy of feedback-based state estimates.

Our study design also included another key distinction from prior work. In bimodal trials of prior studies, individuals were encouraged to integrate visual and proprioceptive information and report a single unified percept (e.g., Block and Bastian, 2011, 2010; Block and Liu, 2023; Reuschel et al., 2010; van Beers et al., 1999, 1996). These prior studies often presented surreptitious sensory mismatches with the intent that people should combine the two senses in some weighted manner to come up with an integrated position percept (Block and Bastian, 2010; 2011; Reuschel et al., 2010). Their approach enabled an assessment of the weights applied to vision and proprioception during sensory integration, but it may have led to the assumption that multisensory integration automatically occurs when deriving a state estimate. In our study, participants were explicitly made aware of the discrepancy between visual and proprioceptive cues, and they were asked to ignore one sensory modality (i.e., deliberately shift the degree to which they relied on one modality versus the other). Thus, our design permitted the assessment of whether multisensory integration is obligatory, but it did not allow us to assess the weights assigned to each modality. Overall, our findings suggest that while multisensory integration may be automatic when the limb is stationary, people can easily ignore the information coming from a second, discrepant modality while moving.

## Conclusion

In summary, we have observed differences in the ability to ignore conflicting sensory information when reporting perceived position versus movement. Our computational model suggests that these differences arise because sensory delays cause bimodal state estimates to be more unreliable at higher movement speeds. Such a pattern of results is consistent with the idea that feedback-based estimates of the current limb state are computed differently when stationary and when moving.

## Acknowledgements

This work is supported by NIH grant R01 NS115862 awarded to ALW, and pilot project funding from the Moss Rehabilitation Research Institute Peer Review Committee awarded to AST and ALW. The authors declare no competing financial interests.

## Author conthributions

ALW and AST conceived and designed the research, interpreted the results of experiments, and edited and revised the manuscript. ALW additionally programmed the experimental tasks, analyzed the data, prepared figures, and drafted the manuscript. LC programmed the experimental tasks, performed the experiments, and assisted with data analysis. AE performed the experiments, and assisted with data analysis.

## Notes

https://osf.io/wt6hz

### Competing Interest Statement

The authors have declared no competing interest.

### Summary of Updates

This revision includes 2 additional control experiments to replicate and generalize our findings from Experiment 1. It has also been reorganized to improve clarity.

